# TLR7-independent control of retroviral infection

**DOI:** 10.64898/2026.04.28.721481

**Authors:** Robert Z. Zhang, Lia Robben, Ashley Willey, Sandra Umana, Vincent Mele, Brady Callahan, Tatyana Golovkina, Melissa Kane

## Abstract

Development of effective immune responses against any pathogen requires efficient activation of the innate immune system followed by induction of an appropriate adaptive immune response using specific signaling pathways tailored to each infection. The sole innate immune receptor implicated in the induction of adaptive immunity to retroviral infection is Toll like receptor 7 (TLR7). We have found that germinal center responses, neutralizing antibody production, and clearance of murine leukemia virus (MLV) infection in BALB.J and C57BL/6N (B6N) mice occurs in the absence of TLR7 signaling. This suggests that a previously unknown alternative sensing pathway exists for the activation of protective immune responses upon retroviral infection. Genetic crosses indicate that the ability to control retroviral infection in the absence of TLR7 signaling is determined by the same recessive mechanism in both B6N and BALB.J mice, suggesting that TLR7-independent responses do not result from a gain-of-function of an alternative pattern recognition receptor. Additionally, we observed that TLR7-deficient BALB.J mice produce neutralizing antibodies against mouse mammary tumor virus (MMTV) infection, indicating that this alternative sensing pathway is active against retroviruses of multiple genera. Finally, we determined that the alternative sensing pathway is also independent of both MyD88 and STING signaling. The ability of mice of two genetic backgrounds to control retroviral infection in the absence of TLR7 signaling provides a valuable tool for the identification of a novel mechanism of retrovirus control. The dissection of this pathway has the potential to alter our understanding of the requirements for the stimulation of antigen-specific neutralizing immunity.

**Significance Statement:** Innate immune sensors are required for induction of pathogen-specific immune responses, and the development of novel vaccines requires an understanding of the basic mechanisms by which the immune system detects and responds to pathogens. While multiple innate immune receptors have been implicated in the sensing of retroviral infection, only Toll-like receptor 7 (TLR7) has been found to upregulate adaptive immune responses. Here, we demonstrate that an additional TLR7-independent mechanism for the activation of antiretroviral antibody responses is present in inbred mice from two genetic backgrounds. This finding has implications for the selection of mouse models for the study of antiviral immune responses and has the potential to alter our understanding of the requirements for the stimulation of antigen-specific neutralizing immunity.

## INTRODUCTION

Resistance and sensitivity to viral infections depends greatly upon the genetic make-up of the host. Model organisms such as inbred mice have proved to be invaluable for the investigation of the pathways underlying protective immune responses due to the inability to perform genetic manipulations in humans. In this regard, mouse models of murine retroviruses, such as murine leukemia virus (MLV) and mouse mammary tumor virus (MMTV) have provided essential insights into the molecular mechanisms underlying anti-viral immune responses. MLV is transmitted as an exogenous virus through the blood and milk or as an endogenous stably integrated provirus and primarily infects cells of lymphoid origin (1). Susceptible mice develop severe splenomegaly and succumb to leukemia. MMTV is also transmitted via both exogenous and endogenous routes. Exogenous MMTV is passed through the milk and initially targets lymphoid cells in the gut which then spread the virus to the mammary glands, leading to tumor development (2).

A required step in the development of a pathogen-specific protective immune response is the recognition of pathogen-associated molecular patterns (PAMPs) by pattern recognition receptors (PRRs) (3). PAMPs represent highly conserved microbial molecular structures that are not found in the host cells or in the compartment of infected cells in which the pathogen replicates. Bacterial pathogens are detected by Toll-like receptors (TLRs), which recognize bacterial lipids, peptidoglycans, or proteins that are foreign to eukaryotic cells (4). Unlike bacterial cell surfaces, viral exteriors lack specific structures that can distinguish them from the surfaces of eukaryotic cells. Consequently, viral recognition occurs through cytosolic or endocytic PRRs that detect virally produced replication intermediates (e.g., various forms of nucleic acids) (5, 6), or through inflammasomes, which detect the activities of some virally encoded proteins (7). Mice from retrovirus-susceptible strains, such as BALB/c mice, sense retroviral pathogens, as indicated by the fact that they initiate an antiretroviral response. However, this response is not long-lasting and is unsuccessful in controlling virus replication (8, 9), which is most likely due to the numerous mechanisms of immune evasion employed by retroviruses (10-14). In contrast, pathogen detection in resistant mice translates into a robust, long-lasting, and virus-neutralizing immune response comprised of both T and B cell responses (8, 10, 15, 16). Previous work by us and others has determined that virus sensing by the endosomal PRR, TLR7 is required for the stimulation of neutralizing antiviral Ab production in retrovirus-resistant mice from two genetic backgrounds, C57BL/6 (B6) and I/LnJ (16, 17).

B6 mice infected with MLV eliminate the virus through both humoral and cell-mediated responses. Antiretroviral antibody (Ab) production in B6 mice is mediated by a single dominant locus, recovery from Friend virus 3 (*Rfv3*), mapped to chromosome 15. *Rfv3* is encoded by the deoxycytidine deaminase *Apobec3* (18) and controls antiviral Ab production via an unknown mechanism. Although B6 mice are resistant to MLV, they are susceptible to MMTV (16). In contrast, mice of the I/LnJ strain control *both* MMTV and MLV by producing virus-neutralizing Abs and cytotoxic T cell responses (8, 19). Retroviral resistance in I/LnJ mice is controlled by a single recessive locus, virus infectivity controller 1 (*vic1*) encoded by the non-classical major histocompatibility class II (MHC-II)-like gene, *H2-Ob* (*HLA-DOB* in humans) (16, 19, 20). BALB/cJ mice congenic for the H2^j^ haplotype [BALB/c-*H2*^*j*^ (BALB.J)] containing the resistant *vic1* allele produce virus-neutralizing Abs upon infection with MLV or MMTV (20, 21).

To date, TLR7 is the only retroviral sensor demonstrated to play a role in the activation of adaptive antiviral immune responses. We report here that production of neutralizing antiretroviral Ab production in BALB.J mice and the B6 substrain, C57BL/6N does not require TLR7 signaling. TLR7-independent antiretroviral immunity is controlled by a recessive genetic mechanism and signals through a MyD88- and stimulator of IFN genes (STING)-independent pathway.

## RESULTS

### Requirement of TLR7 for control of retroviral infection is dependent on genetic background

We tested the requirement for TLR7 in control of retroviral infection in BALB.J mice (**Fig. S1A**) and two C57BL/6 substrains, B6N.TLR7^KO^ mice generated by Richard Flavell [C57BL/6Ncr substrain and available at the Jackson Laboratory on a mixed B6N/B6J background (22)], and B6Jcl.TLR7^KO^ mice generated by Shizuo Akira [C57BL/6Jcl (CLEA) substrain and utilized in our previous report (16, 23)]. Mice were infected with Rauscher-like-MLV (RL-MLV) (24, 25) and viral titers in the spleen were determined at eight weeks post-infection (**Fig. 1A**). While B6Jcl.TLR7^KO^ mice failed to control RL-MLV infection as expected, surprisingly, both BALB.J and B6N mice cleared the infection even in the absence of TLR7. We next tested their sera for the presence of antiviral Abs and found that TLR7-deficient BALB.J and B6N mice produced antiviral Abs that neutralized RL-MLV infection, while TLR7-deficient B6Jcl mice did not (**Figs. 1B-C**). This indicates that TLR7 is not the only innate immune sensor capable of inducing humoral immunity against retroviral infection, and that host genetic background determines the capacity to produce TLR7-independent antiretroviral Abs. Since TLR signaling affects class switch recombination (CSR) (26), we next assessed the isotype specificity of anti-MLV Abs from infected WT and TLR7-deficient BALB.J and B6N mice (**Figs. 1D** and **S2**). Production of MLV-specific IgG2a/c Abs was dramatically reduced in the absence of TLR7-signaling, while IgG1-specific Abs against MLV were not affected by TLR7-deficiency in mice from either background. Isotype distribution was also determined by the genetic background, as production of IgG1 was higher in BALB.J mice than in B6N mice, while the inverse was true for IgG2b-specific Abs against RL-MLV, which were significantly lower in BALB.J mice.

**Figure 1.**
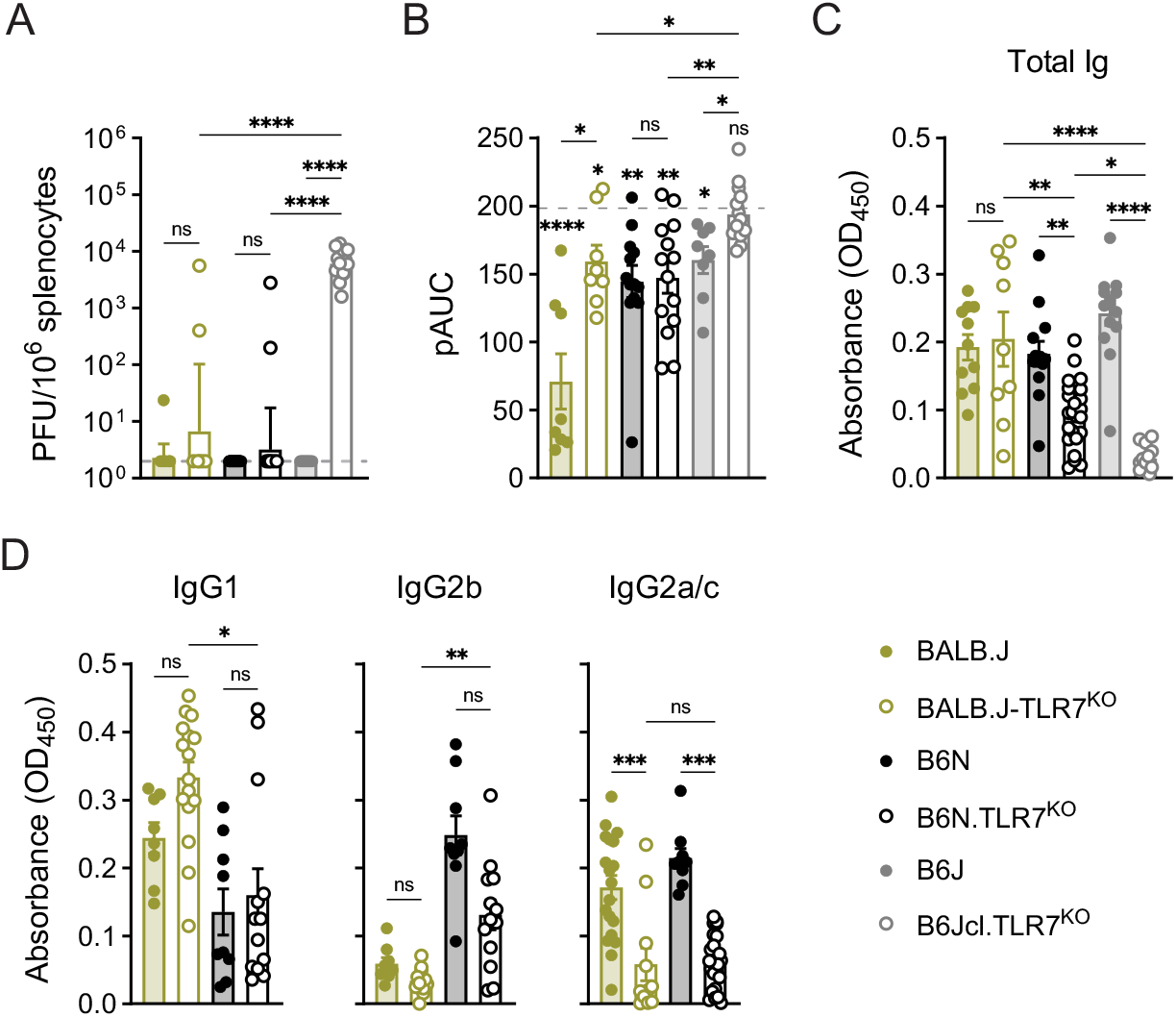
Production of TLR7-independent neutralizing antibodies upon retroviral infection is strain specific. Mice of the indicated genotypes were infected with RL-MLV and analyzed 8 weeks post infection. (A) Splenocytes were isolated and subject to an XC plaque assay, (B) sera was collected, serially diluted and incubated with ecotropic RL-MLV-mNeonGreen reporter virus before adding to SC-1 cells. Neutralization capacity was quantified by determining the partial area under the curve (pAUC). Grey dotted line represents the average pAUC of naïve sera. Statistics above columns represent comparisons against naïve sera. (C) Sera was monitored for total Igs (D) IgG1, IgG2b, and IgG2a antibodies against RL-MLV by enzyme-linked immunosorbent assay (ELISA). *For each statistical comparison a nonparametric Kruskal-Wallis test was applied and corresponding significance values are indicated for each graph. ns, not significant; *, p<0*.*05; **, p<0*.*01; ****, p<0*.*0001*

### Requirement for TLR7 for B and T cell responses upon retroviral infection is dependent on genetic background

We next assessed how host genetic background affects the requirement for TLR7 signaling in germinal center (GC) responses. We conducted a time course to assess the kinetics of humoral responses upon RL-MLV infection and found that the peak GC response was at three weeks post-infection in B6 substrains and at four weeks post-infection in BALB.J mice (**Figs. S3A** and **S4**). While TLR7 signaling was required for the induction of GC responses in B6Jcl mice, WT and TLR7-knockout B6N and BALB.J mice exhibited GC responses of similar magnitude and kinetics (**Figs. 2A and S4**). Similar to our findings regarding isotype specificity of antiviral Abs (**Fig. 1D**), the proportion of B cells that were IgG2a/c positive at the peak of the GC response was reduced in BALB.J and B6N TLR7-deficient mice (**Fig. 2B**). High levels of GC B cells underwent CSR to the IgG1 isotype in both MLV-infected WT and TLR7-knockout BALB.J mice, while TLR7-deficiency increased CSR recombination to IgG1 in B6N mice. These results indicate that the requirement for TLR7-signaling in antiretroviral GC responses is dependent upon host genetic background, and that TLR7 is required for efficient CSR to the IgG2a/c isotype upon retroviral infection.

**Figure 2.**
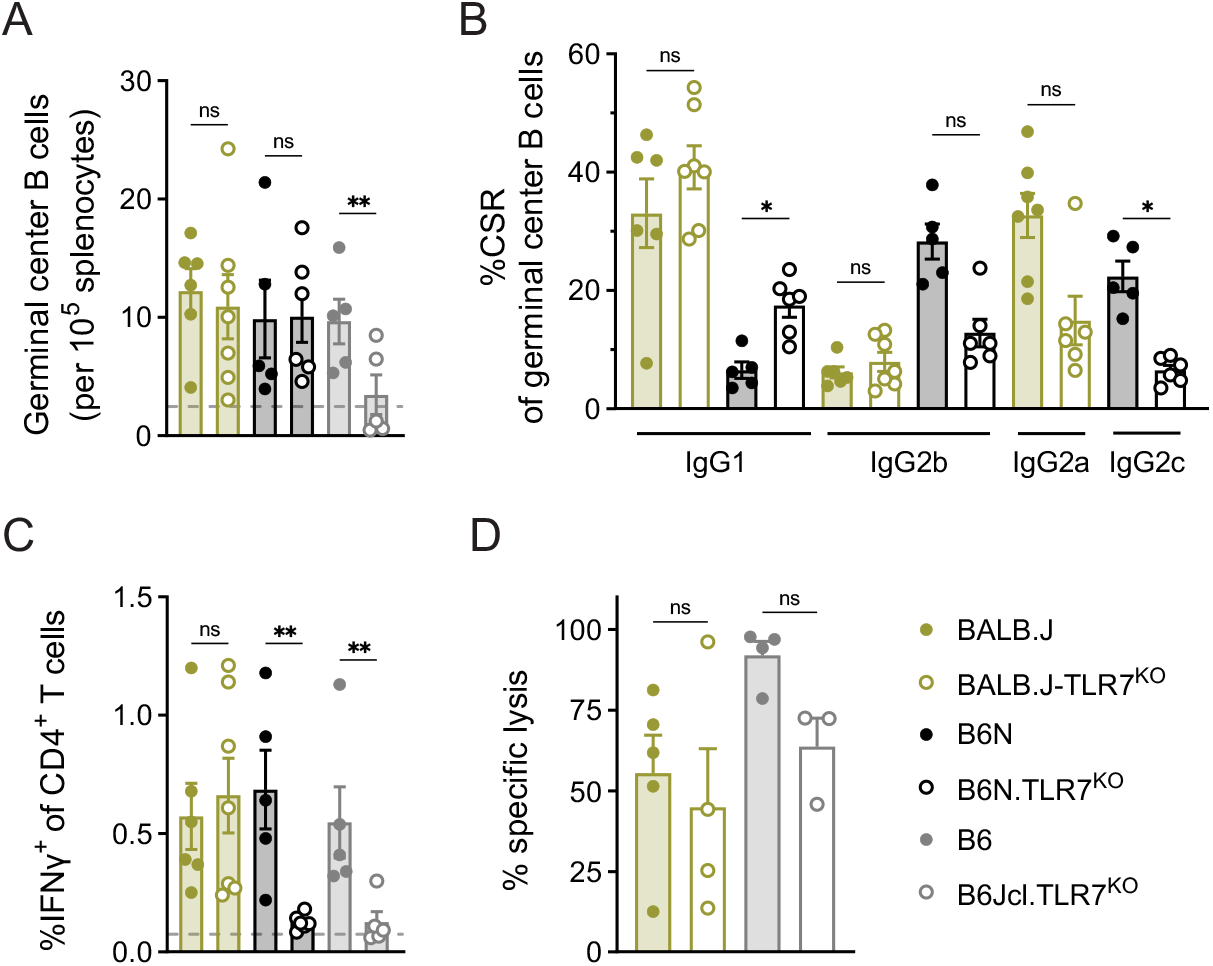
Specific B and T cell responses upon retroviral infection are dependent on genetic background. Mice of the indicated genotypes were infected with RL-MLV and splenocytes were isolated and analyzed by flow cytometry at the peak of germinal center (GC) response (**Fig. S4**). (A) Proportion of GC B cells (Live^+^B220^+^CD95^hi^CD38^lo^). (B) Frequency of GC B cells that were IgM^-^IgG1^+^, IgM^-^ IgG2b^+^, IgM^-^IgG2a^+^, or IgM^-^IgG2c^+^. (C) Frequency of CD4^+^ T cells (Live^+^CD3^+^CD8^-^CD4^+^) that were IFNγ^+^ (D) Mice of the indicated genotypes were infected with RL-MLV. Donor splenocytes were incubated with either MLV GagL peptide, or DMSO. GagL loaded splenocytes were incubated with 5uM CFSE, while non-loaded splenocytes were incubated with 0.5uM CFSE. The CFSE-labeled populations are mixed at a 1:1 ratio and injected intravenously into infected mice. Splenocytes from infected mice were isolated 36 hours post injection and analyzed for CFSE expression by FACS and the percentage of specific lysis of peptide-loaded cells was calculated. *For each statistical comparison a nonparametric Kruskal-Wallis test was applied and corresponding significance values are indicated for each graph*., *not significant; *, p<0*.*05; **, p<0*.*01; ****, p<0*.*0001*

Since GC responses are regulated by CD4 T cells, we also investigated the requirement for TLR7 signaling in CD4 T cell responses in MLV-infected mice (**Figs. 2C, S3B**, and **S5A-D**). IFNγ expression in CD4 T cells was upregulated in WT mice of all three backgrounds following MLV infection, while TLR7-signaling was required for IFNγ expression in CD4 T cells from B6N and B6Jcl mice, but not BALB.J mice (**Fig. 2C**). We also evaluated the proportion of various CD4 T cell subsets upon MLV infection and found that the numbers of T follicular helper (Tfh) and T helper 1 (Th1) cells were unaffected by TLR7-deficiency in mice of all three backgrounds (**Fig. S5A-B**), while a higher proportion of T follicular regulatory (Tfr) cells was found in *Tlr7*^KO^ versus WT B6N and B6Jcl, but not BALB.J mice (**Fig. S5C**). Collectively, these data suggest that while the role of TLR7 in activation of both CD4 T cell and B cell responses to MLV infection is dependent on genetic background, the requirement of TLR7-signaling in CD4 T cell responses is independent of whether it is necessary for antiretroviral Ab production.

Finally, we investigated whether genetic differences affected whether TLR7 signaling is required for CD8 T cell responses to RL-MLV infection (**Figs. 2D** and **S5E-F**). We found that IFNγ expression in CD8 T cells upon RL-MLV infection was not impacted by TLR7 signaling, similar to previous findings by Ed Browne using FV infection (17). We next assessed cytotoxic T cell activity by measuring the ability of infected animals to lyse cells presenting an immunodominant H-2D^b^-restricted GagL peptide (27, 28). WT and TLR7-deficient BALB.J and B6Jcl mice efficiently lysed GagL peptide-loaded cells (**Fig. 2D**), indicating that anti-MLV CTL responses are TLR7-independent. In our previous report, we found that antiviral Ab production and clearance of RL-MLV infection was independent of TLR3 and TLR9 signaling in B6J and I/LnJ mice (16), here we observed that anti-MLV CTL responses were also independent of TLR3 and TLR9 signaling in both backgrounds (**Fig. S5F**). These data indicate that (at least individual) nucleic acid sensing TLRs are dispensable for the activation of CD8 T cell responses upon MLV infection and that the genetic determinants underlying the capacity to produce TLR7-independent antiviral Ab responses do not affect CTL responses.

### TLR7-independent antiretroviral antibody production is controlled by a recessive mechanism

Since B6N.TLR7^KO^ mice are on a mixed B6N/B6J background, and the B6Jcl.TLR7^KO^ mice in this investigation have been bred by independent laboratories for an unknown number of generations, we assessed their genetic divergence via miniMUGA genotyping array (29, 30). This analysis revealed 120 divergent SNPs throughout their genomes, with groups of divergent SNPs (fewer than 4Mb apart) enriched on Chromosomes 7 and 10 and were not linked to *Tlr7* or the FV resistance gene *Rfv3* (**Fig. S6**). Interestingly, this is fewer than the number of SNPs that distinguish C57BL6/J and C57BL6N/J or C57BL/6NCrl (193 and 170, respectively).

To determine whether the capacity for TLR7-independent retroviral control is inherited in a dominant or recessive fashion, we crossed *Tlr7*^KO^ B6Jcl mice to both *Tlr7*^KO^ B6N and *Tlr7*^KO^ BALB.J mice (**Fig. 3**). F_1_ mice generated from these crosses were infected with RL-MLV and phenotyped at eight weeks post-infection. F_1_ mice from both crosses failed to clear the infection (**Fig. 3A**), and although some F_1_ animals produced anti-MLV Abs (**Fig. 3B**), their sera did not neutralize RL-MLV (**Fig. 3C**), indicating that TLR7-independent control of retroviral infection in BALB.J and B6N mice is inherited in a recessive fashion. Additionally, the phenotype of F_1_ animals was independent of the direction of the cross, indicating that the mechanism is not sex-linked and is not affected by the 129 substrain from which the ES cells were derived for the generation of the *Tlr7*^KO^ B6 mice. We next sought to investigate whether TLR7-independent antiretroviral Ab production in BALB.J and B6N mice is controlled by the same genetic mechanism. To test this, we crossed *Tlr7*^KO^ B6N to *Tlr7*^KO^ BALB.J mice. Since TLR7-independent control of MLV is a recessive trait, we reasoned that F_1_ animals would produce neutralizing Abs and control infection only if the phenotype is controlled by the same gene (or pathway). F_1_ mice generated from these crosses were infected with MLV and assessed for their capacity to control infection (**Fig. 3**). Sera from these F_1_ progeny neutralized RL-MLV (**Fig. 3C**), and the majority (16 of 23) either eliminated the virus or had fewer than 1000 PFU per 10^6^ splenocytes, suggesting that the same genetic mechanism controls TLR7-independent antiretroviral immunity in BALB.J and B6N mice.

**Figure 3.**
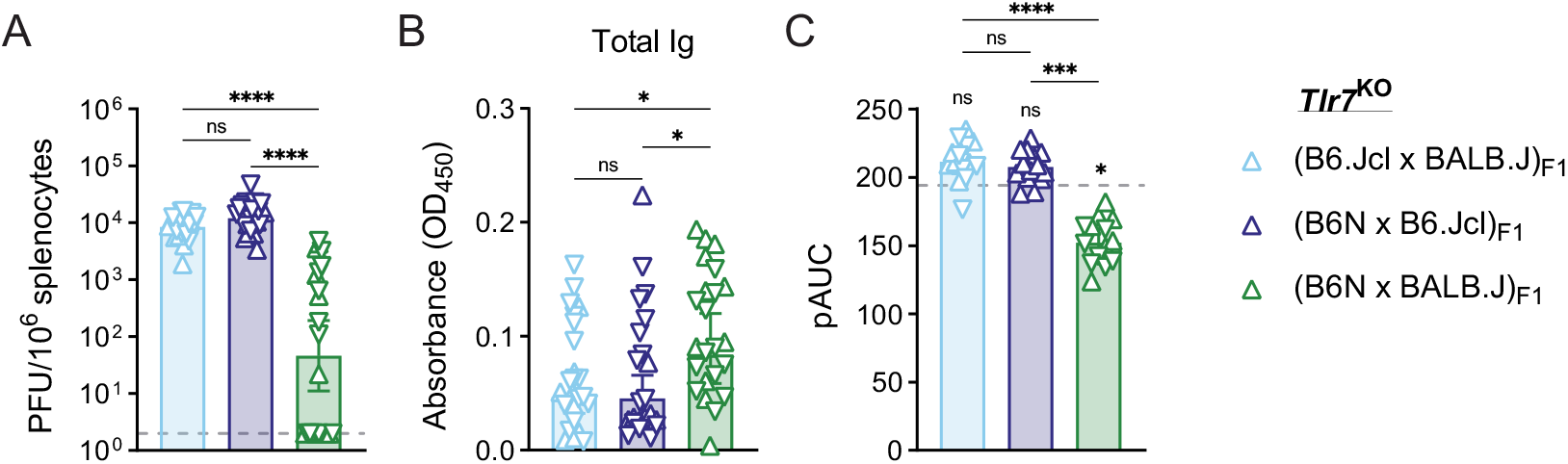
TLR7-independent antiretroviral antibody production is controlled by a recessive mechanism. TLR7-deficient F_1_ mice of the following genotypes, light blue triangle (BALB.J x B6.Jcl) inverted light blue triangle (B6.Jcl x BALB.J), dark blue triangle (B6N x B6.Jcl), inverted dark blue triangle (B6.Jcl x B6N), green triangle (B6N x BALB.J), and inverted green triangle (BALB.J x B6N) were infected with RL-MLV and analyzed at 8 weeks post infection. (A) Splenocytes from infected mice were isolated and subjected to an XC plaque assay (B) Sera from infected mice was monitored for total Igs against RL-MLV by enzyme-linked immunosorbent assay (ELISA). (C) Sera from infected mice was serially diluted and incubated with ecotropic RL-MLV-mNeonGreen reporter virus before adding to SC-1 cells. Neutralization capacity was quantified by determining the partial area under the curve (pAUC). Grey dotted line represents the average pAUC of naïve sera. Statistics above columns represent comparisons against naïve sera. *For each statistical comparison a nonparametric Kruskal-Wallis test was applied and corresponding significance values are indicated for each graph. ns, not significant; *, p<0*.*05; ****, p<0*.*0001*

### Control of retroviral infection in BALB.J mice in the absence of MyD88 or STING signaling

We next sought to identify the innate immune signaling pathway responsible for activation of TLR7-independent antiretroviral immunity. Since variation between C57BL/6 substrains affects the requirement for TLR7 in anti-MLV responses and the genetic determinants that underlie these differences are unknown, we utilized BALB.J mice to investigate the TLR7-independent signaling pathway. Multiple TLRs in addition to TLR7 have been implicated in sensing retroviral infection (31). The adaptor molecule MyD88 is required for most TLR signaling, except for TLR3 which signals through the adaptor Ticam1, and TLR4 which can signal through either MyD88 or Ticam1 (6). We infected *Myd88*^KO^ BALB.J mice with RL-MLV and monitored them for antiviral Ab production and viral titers in the spleen (**Fig. 4**). We found MyD88-deficient BALB.J retained the capacity to produce anti-MLV neutralizing Abs (**Fig. 4B**), although similar to our findings in *Tlr7*^KO^ BALB.J mice (**Fig. 1C**), sera from *Myd88*^KO^ mice exhibited reduced neutralization potency as compared to WT mice (**Fig. 4C**), and three of seven MyD88-deficient animals did not control viral replication (**Fig. 4A**). Loss of MyD88 signaling also affected isotype distribution of anti-MLV similar to TLR7-deficiency, as *Myd88*^KO^ mice produced lower titers of IgG2a-, but not IgG1- or IgG2b-specific Abs against MLV (**Fig. 4D**). These data suggest that while full protection against MLV infection requires MyD88-signaling, the TLR7-independent pathway for antiretroviral Ab production is MyD88-independent.

**Figure 4.**
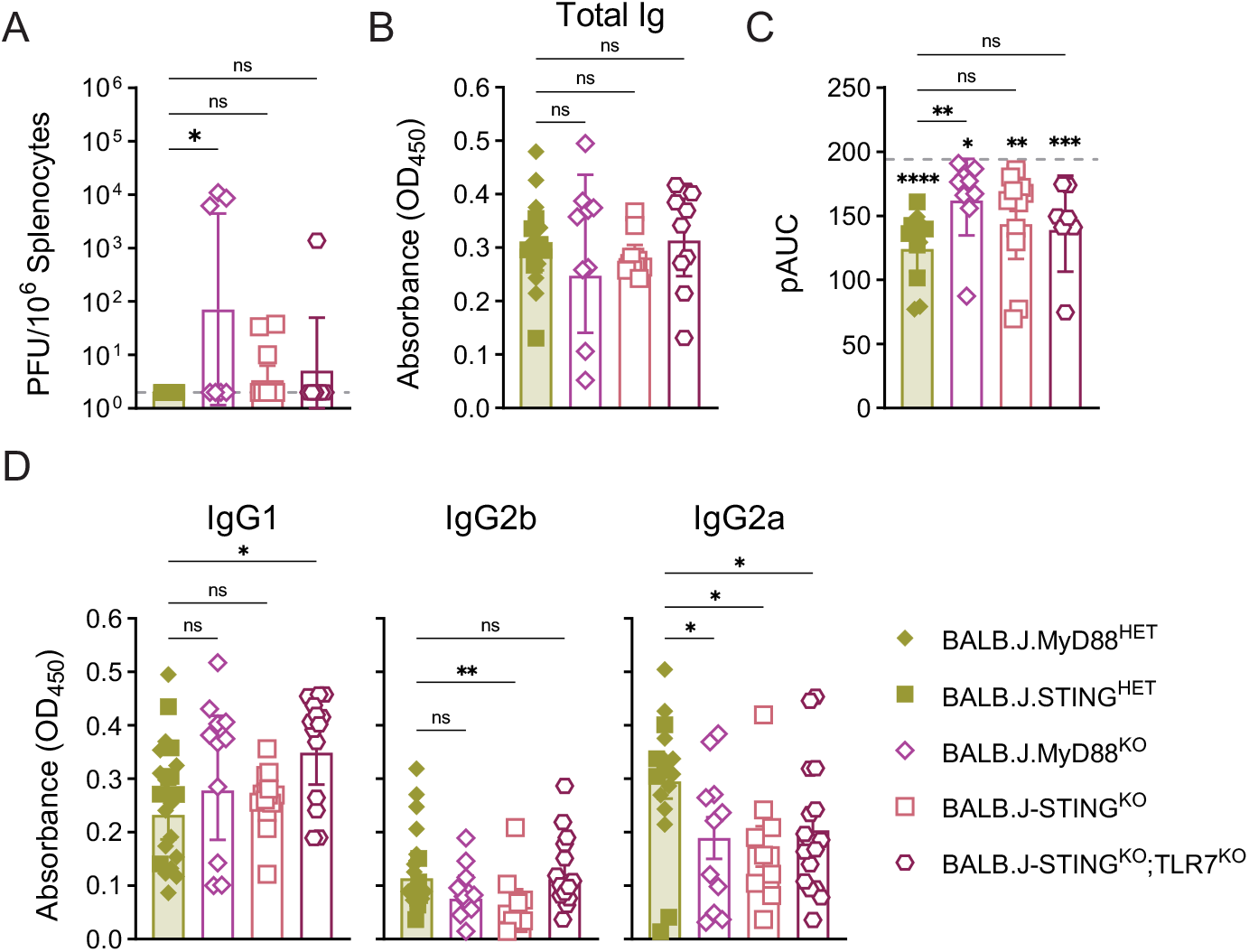
TLR7-independent protective immune responses are not controlled by STING or MyD88 signaling. Mice of the indicated genotypes were infected with RL-MLV and analyzed 8 weeks post infection. (A) Splenocytes from infected mice were isolated and subjected to an XC plaque assay (B) Sera was monitored for total Igs against RL-MLV by enzyme-linked immunosorbent assay (ELISA). (C) Sera from infected mice was serially diluted and incubated with ecotropic RL-MLV-mNeonGreen reporter virus before adding to SC-1 cells. Neutralization capacity was quantified by determining the partial area under the curve (pAUC). Grey dotted line represents the average pAUC of naïve sera. Statistics above columns represent comparisons against naïve sera. *For each statistical comparison a nonparametric Kruskal-Wallis test was applied and corresponding significance values are indicated for each graph. ns, not significant; *, p<0*.*001; **, p<0*.*01*, ***, p<0.0001; ****, p<0.0001

In addition to TLR7-mediated sensing of retroviral RNA, reverse transcripts are sensed by cellular DNA sensors such as cyclic GMP-AMP synthase (cGAS), DExD/H-box helicase 41 (DDX41), and the absent in melanoma 2 (AIM2)-like receptors (ALRs) interferon-induced 16 (IFI16) in humans and IFI203 in mice (32-37). These sensors signal through the common adaptor, stimulator of IFN genes (STING), which results in the expression of type I IFNs and antiretroviral restriction factors such as APOBEC3 (32, 38). This STING-dependent pathway has been shown to be involved in the control of viral load in B6 mice (32, 36), but has not been implicated in the activation of adaptive immune responses. We generated *Sting*^KO^ mice on the BALB.J background (**Fig. S1B-C**) and assessed antiviral Ab production and viral titers in the spleen in MLV-infected *Sting*^KO^ and *Sting*^KO^;*Tlr7*^KO^ mice (**Fig. 4**). STING-deficient mice produced neutralizing anti-MLV Abs and controlled the infection with or without TLR7-deficiency (**Fig. 4A-C**) indicating that TLR7-independent antiretroviral immunity is independent of the STING pathway. Interestingly, we also found that *Sting*^KO^ mice produced lower titers of IgG2a-specific anti-MLV Abs, and *Sting*^KO^;*Tlr7*^KO^ mice had elevated titers of IgG1-specific anti-MLV Abs (**Fig. 4D**), suggesting that cytokine production downstream of both MyD88- and STING-dependent pathways affects CSR upon retroviral infection.

### TLR7-independent production of neutralizing Abs against MMTV

In our previous report, we found that like MLV, production of Abs against MMTV in I/LnJ mice and B6-*H2*^*j*^ mice is TLR7-dependent (16). To determine whether the TLR7-independent antiretroviral Ab production in BALB.J mice is restricted to control of MLV infection, we infected WT and *Tlr7*^KO^ BALB.J mice with MMTV and monitored them for antiviral Ab production (**Figs. 5** and **S7**). We found that BALB.J mice produced neutralizing anti-MMTV Abs in the absence of TLR7-signaling, albeit with lower potency than WT mice (**Fig. 5A**). We also examined the isotype distribution of MMTV-specific Abs and TLR7-deficiency reduced CSR to IgG2a and IgG1, but not IgG2b-specific Abs (**Fig. 5B**). Thus, although TLR7 signaling affects isotype distribution of antiviral Abs, it is not required for the production of neutralizing Abs against MMTV. Furthermore, the TLR7-independent pathway for stimulation of antiretroviral immunity is not limited to MLV infection and can control retroviruses from distinct genera.

**Figure 5.**
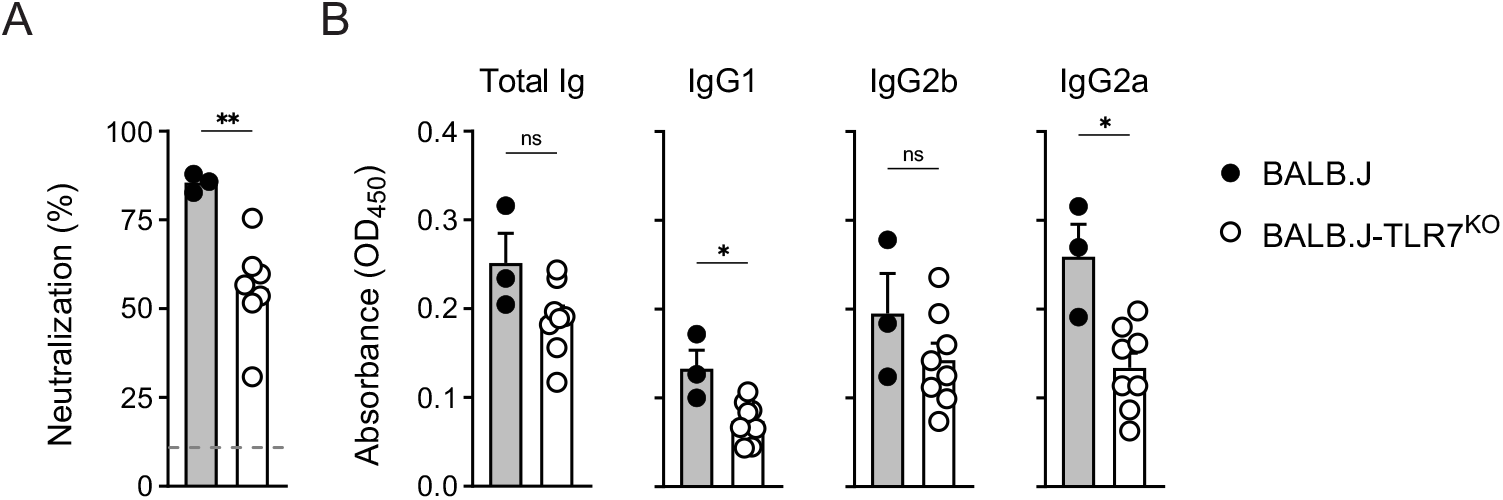
MMTV infection stimulates the production of TLR7-independent neutralizing antibodies in BALB.J mice. BALB.J (closed circles) and TLR7-deficient BALB.J mice (open circles) were infected with MMTV(LA). (A) Sera from infected mice was collected 8 weeks post infection and mixed with purified MMTV virions and tested for neutralization of MMTV in BALB/cJ mice. Sera from a naïve mouse was used as a control and is represented by the grey dotted line. (B) Sera was incubated with monitored for total Igs, IgG1, IgG2b, IgG2a against MMTV virions by enzyme-linked immunosorbent assay (ELISA). *For each statistical comparison a nonparametric Mann-Whitney test was applied, and corresponding significance values are indicated for each graph. ns, not significant; *, p<0*.*001; **, p<0*.*01*,

## DISCUSSION

Here, we provide evidence for a previously unknown sensing pathway for the activation of protective adaptive immune responses against retroviral infection. Whereas previous work by us and others indicated that TLR7 is the sole PRR responsible for the activation of humoral immunity upon retroviral infection, we identified two strains of mice capable of GC formation and neutralizing antiretroviral Ab production in the absence of TLR7 signaling. This finding indicates that host genetic variation affects the requirement for specific innate immune receptors in the activation of adaptive immune responses and further highlights the importance of utilization of mice from multiple genetic backgrounds to determine the role of a given pathway in antiviral immunity whenever possible.

TLR7-independent antiretroviral immunity is controlled by a recessive genetic mechanism (**Fig 3**), suggesting that it does not involve a gain of function of an alternative sensor in B6N and BALB.J mice. Rather, our findings suggest that either a suppressive or negative regulatory pathway is activated in strains that inherit the dominant allele and require TLR7 signaling for antiretroviral immunity, or that activation of TLR7-independent neutralizing immunity is dosage-sensitive, requiring two copies of the resistant allele(s). These could include a cytokine or upstream regulator of B cell responses, an immune receptor or signaling adaptor, or a noncoding element that affects expression of another gene(s) that directs antiviral immunity. Complementation of the BALB.J and B6N alleles indicates that their capacity to produce antiviral Abs in the absence of TLR7 is controlled by the same gene or a similar pathway. Since the phenotype of the F_1_ progeny is not fully penetrant, it is likely that they do not inherit identical alleles, however the identification of two strains with a similar mechanism will aid in future investigation of the genetic basis for this novel pathway.

Our results add to the growing list of phenotypic differences among C57BL/6 substrains that both provides an opportunity for novel gene discovery and creates confounding effects when these differences are not rigorously accounted for (39, 40). Notably, the *Tlr7*^-/-^ B6N mice used in this investigation are the same strain as those reported by Ed Browne in 2011 to require TLR7 signaling for anti-retroviral GC responses upon Friend virus (FV) infection (17). While our data regarding the role of TLR7 in activation of CD4 and CD8 T cell responses upon MLV infection in B6 mice are in agreement with his report, our findings regarding the role of TLR7 in antiretroviral Ab production are contradictory. Importantly, Browne did find that early upregulation of CD69 and CD86 expression on B cells was independent of TLR7 signaling, suggesting that the TLR7-independent pathway was activated upon FV infection, but the humoral response failed to progress (17). Since the mice sent to the Jackson Laboratory Repository were on a mixed B6N/B6J background and SNP analysis for genetic quality control was not conducted until 2021 (41), it is possible that genetic divergence between the mice in the Browne report and those used in our investigation explain this discrepancy, especially given the differences between TLR7-deficient B6Jcl and B6N mice that we report here. Alternatively, differences between the biology of FV and RL-MLV complexes could affect innate and/or adaptive immune signaling pathways. For example, FV contains a replication-defective spleen focus forming virus (SFFV) component that promotes pathogenesis (1), while RL-MLV does not (24), thus the discrepancy could also be explained by inhibition of GC responses in TLR7-deficient mice by SFFV.

Cytokine-induced CSR resulting in the production of specific Ab isotypes is an essential step in the development of effective antiviral humoral responses. This process involves B cell receptor stimulation, TLR and CD40 signaling, as well as interactions with helper T cells and the activation of isotype-specific promoters by cytokines (42, 43). The production or enrichment of specific isotypes following infection influences the effectiveness of antiviral immune responses in both mice and humans due to functional differences between different Ab isotypes, including the capacity to activate complement and affinity for Fcγ receptors (44). In mice, IgG2a/c Abs are the predominant isotype elicited upon with a variety of viruses and parasites including retroviruses (8, 19, 45-48). Additionally, numerous reports have indicated that TLR signaling affects GC responses and IgG2a/c CSR, in both a B cell-intrinsic and -extrinsic manner (49-53). In agreement with these earlier investigations, we found that efficient CSR to IgG2a/c upon MLV or MMTV infection requires TLR7/MyD88 signaling (**Figs 1C, 2B, 4D**, and **5B**). Surprisingly, we also found that similar to MyD88-deficiency ablation of STING signaling reduced CSR to IgG2a, and that anti-MLV IgG2a titers were not further reduced in STING/TLR7-knockout mice (**Fig. 4D**). These findings suggest that STING signaling affects TLR7-dependent antiretroviral Ab responses. Responsiveness to TLR7 stimulation in B cells is uniquely dependent on type I IFN signaling (54-56). Therefore, STING activation may be affecting the TLR7-dependent pathway for antiretroviral Ab production and CSR via the production of type I IFNs. In addition to differences in the isotype distribution of anti-MLV Abs between B6N and BALB.J mice (**Fig. 1D**) we also found that the isotype profile of antiretroviral Abs was distinct in MLV and MMTV infected BALB.J mice. The subclass hierarchy in MLV-infected mice was IgG1>IgG2a>IgG2b while MMTV infection resulted in IgG2a>IgG2b>IgG1 (**Figs. 1D** and **5B**). This divergent profile suggests that distinct immune pathways activated upon MLV and MMTV infection differentially shape the Th1/Th2 balance of the antiviral Ab response, possibly through infection and/or sensing in distinct cell types. Overall, this investigation demonstrates that the isotype distribution of antiretroviral Abs is affected by TLR7 signaling, host genetic background, and the virus itself.

We eliminated both MyD88- and STING-dependent signaling pathways are involved in the stimulation of TLR7-independent antiretroviral Abs – what then is the alternative innate immune pathway? One possibility is TLR3 which senses dsRNA and signals through a MyD88-independent mechanism (57, 58). Although retroviruses do not produce a dsRNA during their replication, the ssRNA genome can form secondary structures or dimers, which could be sensed in the endosome. TLR3 has also been implicated in control of FV infection, but not via the stimulation of humoral immunity (59). Another possibility is that retroviral infection may be detected by the cytosolic RIG-I like receptor (RLR) RIG-I or MDA5. These sensors are required for type I IFN production against several viral infections, including VSV, influenza virus, murine norovirus 1, and murine hepatitis virus (60). However, a role for the RLRs in activation of adaptive immune responses has not been identified. Both RIG-I and MDA5 recognize RNA targets that are unique to virally infected cells (5′-triphosphate moieties and long double-stranded RNA [dsRNA] molecules, respectively). Since retroviral RNA is produced by cellular RNA polymerases, it was unclear how it would activate either of these sensors until a recent report demonstrated that intron-containing RNA expressed from the integrated HIV-1 provirus activates MDA5 (61). Many viral infections, including VSV and influenza virus, activate the inflammasome, stimulating caspase 1 activation and secretion of interleukin-1β (IL-1β) and IL-18 (62). HIV-1 activation of the NLRP3 inflammasome in monocytic cells has been implicated in neuronal damage (63-65), but this may be caused by cytopathic effects of viral infection rather than detection of viral nucleic acids; furthermore, there is no indication that it is stimulated by MLV or MMTV infection. It is also possible that a yet-to-be-identified sensor initiates TLR7-independent immunity. Identification of the alternative sensor and the retroviral ligand, as well as the genetic mechanisms that determine whether this pathway is functional will be key areas for future investigation. Dissection of this previously unappreciated pathway may have implications for the design of protective vaccines against various viruses.

## MATERIALS and METHODS

Detailed information regarding mice, cell lines, and viruses used is provided in *Materials and Methods* and SI References. The methods for generating and genotyping mice, immunoblot, infection, XC plaque assay, ELISA, neutralization assays, flow cytometry, *in vivo* cytotoxic killing assay, and statistical analysis are described in *Materials and Methods* and SI References. Antibodies and flow cytometry reagents used are listed in *SI Appendix*, Table S1.

## ACKNOWLEDGMENTS

We thank S. Gingras and the transgenic and gene targeting (TGT) core for the generation of *Tlr7*^*-/-*^ and *Sting*^*-/-*^ BALB/c mice and Rebecca Elsner for providing technical advice regarding spectral flow cytometry analysis. Color-blind safe qualitative color schemes from Paul Tol were used for figures (66). This work was supported by grants from the NIH: R21AI178197 (MK), R01CA134667 (TG) and T32 AI049820 (RZ), by the Diane and Cliff Rowe Research Fund (MK).

## Notes

### Competing Interest Statement

The authors have declared no competing interest.

