## Supplementary Information for "TLR7-independent control of retroviral infection"

**This PDF file includes:**

Supplemental Materials and Methods
Figures S1 to S7
Table S1
SI References

### SI Materials and Methods

#### Mice

BALB.J mice were described previously and were bred and maintained at the University of Pittsburgh (1). C57BL/6NJ (stock #005304), C57BL/6J (stock #000664), B6.129S1-*Tlr7*$^{tm1Flv}$/J (stock #008380), and BALB/cJ (stock #000651) were purchased from the Jackson Laboratory. B6.129P2-*Tlr7*$^{tm1Aki}$ were a gift from Thirumala-Devi Kanneganti (St. Jude Children's Research Hospital) with the permission of Dr. Shizuo Akira. MyD88-deficient BALB/c mice were obtained from Mark Shlomchik *via* cryorecovery from sperm at the University of Pittsburgh (2) and subsequently crossed to BALB.J mice to generate MyD88-deficient BALB.J mice. C3H/HeN mice were a gift from Tatyana Golovkina (University of Chicago).

*Tlr7*-deficient BALB/cJ mice were generated using CRISPR/Cas9 technology. The target sequences (*Tlr7*-proximal guide: 5'-ATCTGAGACACCTAATTGGAGGG-3'; *Tlr7*-distal3 guide: 5'-TTCATAGTAAGCCCAAAAGGGGG-3') were previously validated for the generation of conditional knockout mice on the MRL/lpr background (3). One male founder with a complete deletion of exon three was selected (Fig. S1A) and crossed to BALB.J mice. Potential off-target sites with three or fewer mismatches identified by the CRISPOR website (4) were sequenced in $N_1$ and $N_2$ mice, and mice with no mutations at these sites were intercrossed to produce homozygous *Tlr7*-deficient BALB.J mice.

*Sting1*-deficient BALB/cJ mice were generated using CRISPR/Cas9 technology. The target sequence (5'-GTTAAATGTTGCCCACGGGC-3') was previously validated for the generation of targeted mutations on the C57BL/6 background (5). One male founder and one female founder with frameshifts in exon five were selected (Fig. S1B) and crossed to BALB.J mice. Potential off-target sites with three or fewer mismatches identified by the CRISPOR website (4) were sequenced in $N_1$ and $N_2$ mice, and mice with no mutations at these sites were intercrossed to produce homozygous *Sting1*-deficient BALB.J mice. *Sting1*-knockout for both lines was confirmed by immunoblot (Fig. S1C).

All mice were bred and maintained at the University of Pittsburgh. Mice were genotyped from tail biopsies using real time PCR with specific probes designed for each gene (Transnetyx, Cordova, TN). Mice of both sexes were used in equal numbers for each experiment, except for MMTV neutralization experiments which were done in all female mice. All animal experiments were performed in the American Association for the Accreditation of Laboratory Animal Care-accredited, specific-pathogen-free facility Division of Laboratory Animal Resources, University of Pittsburgh School of Medicine. Animal protocols were reviewed and approved by the Institutional Animal Care and Use Committee at The University of Pittsburgh.

The following mice were used for the *in vivo* cytotoxic killing assay and were bred and maintained at the University of Chicago. C57BL/6J (stock #000664), I/LnJ (stock #000674), and B6;129S1-*Tlr3*$^{tm1Flv}$/J (stock #005217) were purchased from The Jackson Laboratory. B6.129P2-*Tlr7*$^{tm1Aki}$ were obtained from Derry Roopenian (The Jackson Laboratory). B6.*Tlr9*-/- mice (6) were obtained from Akiko Iwasaki (Yale University). I/LnJ.*Tlr3*-/-, I/LnJ.*Tlr7*-/-, I/LnJ.*Tlr9*-/-, and I/LnJ.CD8-/- mice have been previously described (7, 8). Animal protocols were reviewed and approved by the Animal Care and Use Committee at the University of Chicago.

**MiniMUGA analysis**

To identify genomic differences between B6.129S1-*Tlr7*$^{tm1Flv}$/J (B6N.TLR7$^{KO}$) and B6.129P2-Tlr7$^{tm1Aki}$ (B6Jcl.TLR7$^{KO}$) strains, we subjected tail biopsies from three mice each (two males and one female) to the MiniMUGA genotyping array (9) (Transnetyx, Cordova, TN). We used Claude.ai to identify divergent SNPs between the two strains (Fig. S6).

**ELISA**

To detect MLV Abs in mouse sera, an enzyme-linked immunosorbent assay (ELISA) was performed as previously described (7, 10). Virions isolated from RL-MLV infected SC-1 cells were treated with 0.1% Triton X-100 and bound to plastic in borate-buffered saline overnight, followed by incubation with mouse serum samples at 4°C for 60 minutes. All sera were used at $2x10^{-2}$ dilutions. Mouse Ig-specific secondary antibodies coupled to horseradish peroxidase

(HRP; Jackson ImmunoResearch) were used to detect antivirus antibodies. Ovalbumin (2%) was used as a blocking reagent. Backgrounds obtained from incubation with secondary antibodies alone were subtracted from the values obtained from sera of infected mice.

**Immunoblot**

To confirm *Sting1* knockout, splenocytes were lysed in NuPage LDS sample buffer (Invitrogen) followed by sonication, separated by electrophoresis on NuPage 4%–12% Bis-Tris, and blotted onto polyvinylidene fluoride (BioRad Laboratories). Membrane was incubated with anti-STING (13647; Cell Signaling 1:1,000) and Anti-TUBA4A (T6074; Millipore Sigma; 1:10,000) primary antibodies followed by incubation with goat anti-rabbit-HRP or goat anti-mouse-HRP secondary antibodies (Jackson ImmunoResearch). SeeBlue Plus2 standard (ThermoFisher) was used. Blot was developed with SuperSignal West Femto Maximum Sensitivity Substrate (ThermoFisher) and imaged on a C-Digit scanner (LI-COR Biosciences).

**Cell lines**

SC-1 embryonic mouse fibroblasts (ATCC CRL-1404) and vir6 cells [SC-1 cells stably infected with the RL-MLV mixture (11)] were maintained in Dulbecco's Modified Eagle Medium [(DMEM) Gibco] with 5% fetal calf serum [(FCS) Gibco] and gentamicin (Gibco). XC (ATCC CCL-165) were maintained in Modified Eagle Medium [MEM (Gibco)] with 10% FCS, sodium pyruvate (Gibco) and gentamicin. HEK293T cells (ATCC CRL-3216) were maintained in DMEM with 10% FCS and gentamicin. Mycoplasma testing (Invivogen) is conducted on all cell lines bi-monthly, and adherent cell lines were maintained in MycoZap Prophylactic (Lonza) at 1:1000 to prevent mycoplasma contamination. Prior to experimental use of cells, medium was changed to remove prophylactic.

**Plasmids and cloning**

The RL-MLV-mNeonGreen reporter construct used for neutralization assays was generated by inserting a P2A sequence followed by the mNeonGreen sequence between the 3' end of *env* and the long terminal repeat (LTR) in the infectious plasmid clone of RL-MLV, pSRL.

Specifically, the sequence between the *Age*I site (located in *env*) and the *EcoR*I site at the end of the LTR, including the P2A and mNeonGreen sequences, was synthesized as a gBlock fragment (Azenta) and subcloned into pSRL using restriction enzymes *Age*I and *EcoR*I (NEB).

**Viruses and infection**

Rauscher-like MuLV (RL-MLV), a mixture consisting of NB-tropic ecotropic and mink lung cell focus-forming viruses, was described previously (11) and was provided by Tatyana Golovkina (University of Chicago). The virus was propagated in vir6 cells. Ecotropic (Eco) viral titers were determined by XC plaque assay (12). Experimental mice were infected *via* intraperitoneal (i.p.) injection with $2\times10^4$ Eco PFUs at 5-8 weeks of age and screened for anti-virus antibodies and plaque forming units 8-10 weeks later.

RL-MLV-mNeonGreen stocks were generated by transfecting HEK293T cells using polyethyleneimine (PolySciences) for 48-72 hours. Harvested supernatant was filtered, and mNeonGreen reporter virus stocks were titered by serial dilution and incubation with SC-1 cells for 48 hours to determine the amount of virus used for experiments.

MMTV(LA), a naturally occurring exogenous virus (13), was used for infection. The virus was propagated in C3H/HeN mice. MMTV(LA) consists of three different exogenous MMTVs, BALB2, BALBLA, and BALB14, with Vβ2-, Vβ6-, and Vβ14-specific super-antigens (SAgs), respectively (13, 14). Mice were infected *via* i.p. injection of 5–8-week-old mice with milk-borne MMTV as previously described (7, 15). Deletion of $CD4^+$/Vβ6 and $CD4^+$/Vβ14 SAg-cognate T cells was used to confirm MMTV infection. FACS analysis of peripheral blood lymphocytes (PBLs) was used to measure deletion rates.

**XC plaque assay**

MLV viral titers were determined by XC plaque assay (12). In brief, isolated splenocytes or vir6 supernatant were incubated with SC-1 cells for 5 days. SC-1 cells were killed by UV-irradiation and overlaid with XC cells for 2 days. Cells were then stained with a mixture of methylene blue

(Thermofisher) and carbol fuchsin (Sigma-Aldrich) in methanol, and localized syncytia were counted.

**Neutralization assays**

Sera from MLV-infected and control mice were tested for their ability to neutralize virus. mNeonGreen reporter RL-MLV virions were incubated with serially diluted heat-inactivated sera from infected mice for one hour at room temperature. Virus/antibody mixtures were added to permissive SC-1 cells. 48 hours later, cells were fixed and infected cells (%mNeonGreen positive) were determined by FACS analysis using an Attune NxT flow cytometer coupled to an autosampler. Neutralization capacity of sera was calculated using the partial area under the curve (16) and normalized to sera from naïve mice.

Sera from MMTV-infected and control mice were tested for their ability to neutralize virus. Each serum diluted at 1/10 with PBS was incubated with purified MMTV(LA) for 2h at room temperature and 25µl of this mixture was injected by hock injection into one hind leg of BALB/cJ mice. Four days after injection, cells isolated from the draining popliteal lymph node were analyzed by FACS for the percentage of $CD4^+/V\beta6^+$ T cells among $CD4^+$ T cells. MMTV-encoded SAg stimulates cognate T cells to proliferate during initial stages of infection (17) and thus, the proliferation of SAg cognate T cells was used as an indicator of virus infectivity (18). The amount of virus prep used in the experiments was first titrated to give an increase from 12% to 25% of SAg-reactive T cells four days after virus injection. Neutralization (%) was calculated as follows: (a-c)-(b-c):(a-c)X100 where a = mean of $CD4^+/V\beta6^+$ (%) in mice injected with virus plus sera from uninfected mice, b = mean of $CD4^+/V\beta6^+$ (%) in experimental mice and c = mean of $CD4^+/V\beta6^+$(%) in uninfected mice.

**Flow cytometry**

Splenocytes were isolated and made into single cell suspensions by mechanical disruption. Red blood cells were lysed with sterile distilled water (Gibco). White blood cells were washed and resuspended in PBS, and live cells were enumerated with trypan blue and Countess 3

Automated Cell Counter (Thermofisher). Cells were pelleted and washed once in PBS before staining. For all stains, two million cells per sample were stained with Zombie Aqua or Zombie NIR Fixable Viability Kit (BioLegend) according to manufacturer's protocol and washed with FACS buffer [PBS (Corning) with 1% bovine serum albumin (Invitrogen), and 0.01% sodium azide)]. Cells were stained with fluorescently conjugated antibodies as labeled in figures for 30 min on ice, then washed with FACS buffer, and fixed with 2% PFA in PBS for 15 minutes on ice. For intracellular stains, cells were fixed/permeabilized with FoxP3 Transcription Factor staining Kit (Invitrogen) according to manufacturer's protocol. After fixation, cells were stained with intracellular stain antibodies as labeled in figures overnight at 4$^o$C. Cells were then washed three times with permeabilization buffer, resuspended in FACS buffer, and data was collected using a Cytek Aurora flow cytometer and analyzed with FlowJo software.

To measure deletion or expansion of CD4$^+$/Vβ6$^+$ PBLs in MMTV-infected mice, leukocytes were recovered from heparinized blood samples by centrifugation through a Ficoll-Hypaque cushion. Cells were resuspended in FACS buffer and stained with CD4-PE Cy7 (Invitrogen) and Vβ6-PE (BD Biosciences). Stained cells were analyzed using an Attune NxT flow cytometer and FlowJo software.

***In vivo* cytotoxic killing assay**

The *in vivo* cytotoxic killing assay was performed as described (19). Briefly, experimental mice were infected with the RL-MLV mixture described above. 12 days later, naïve donor splenocytes were isolated and half were incubated with a modified MLV GagL peptide (Abu-Abu-Leu-Abu-Leu-Thr-Val-Phe-Leu) (20, 21), purchased from GenScript. Peptide-loaded and unloaded cells were then stained with 5μm (CFSE$^{hi}$) and 0.5μm (CFSE$^{lo}$) of CFSE, respectively. Cells were then mixed at a 1:1 ratio, and 2x10$^6$ total cells were intravenously (i.v.) injected *via* tail vein injection into experimental mice. 36-40h later, splenocytes of experimental mice were isolated, and the ratio of CFSE$^{hi}$ and CFSE$^{lo}$ splenocytes was analyzed using an Attune NxT flow cytometer and FlowJo software. Percent specific lysis was calculated as follows:

Ratio in naïve mouse: (%$CFSE^{lo}$)/(%$CFSE^{hi}$)

Ratio in infected mouse: (%$CFSE^{lo}$)/(%$CFSE^{hi}$)

% specific lysis = [1-(naïve ratio/infected ratio)]x100

We validated that the % specific lysis in uninfected mice was zero.

**Statistical analysis**

We use nonparametric Mann-Whitney tests to analyze two-group comparisons. Multi-group comparisons were analyzed by nonparametric Kruskal-Wallis test. All statistical analyses were performed with GraphPad Prism 11, with significance defined as $P < 0.05$.

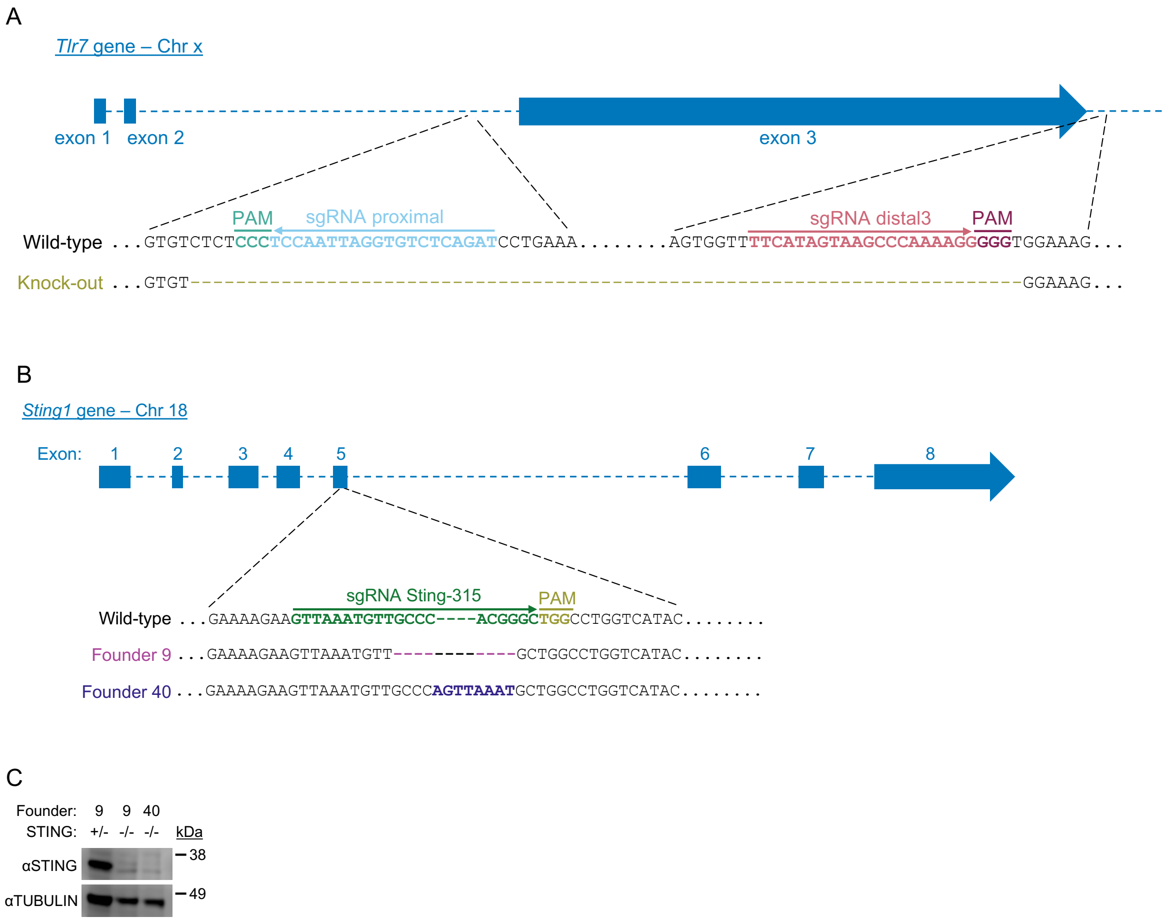


**Supplemental Figure 1. Generation of TLR7-deficient and STING-deficient mice.** (A) Diagram of the *Tlr7* locus, with sgRNA targeting sites in intron 2 and intron 3 indicated. Sequence of wild-type mice at targeted sites, with sgRNA and PAM sites indicated. Dots indicate sequences flanking those shown in detail. Sequences of the locus in selected founder mice with mismatches to wild-type sequence highlighted, deleted sequence indicated by dashes. TLR7-deficient mice lack exon 3, which encodes for amino acids 2-1050 (B) Diagram of the *Sting1* locus, with the sgRNA targeting site in exon 5 indicated. Sequence of the wild-type mice at targeted site, with sgRNA and PAM sites indicated. Dots indicate sequences flanking those shown in detail. Sequences of the locus in selected founder mice 9 and 40 with mismatches to wild-type sequence and insertion highlighted and deleted sequence indicated by dashes. (C) Western blot analysis for STING expression and TUBULIN loading control in splenocytes for mice with the indicated genotypes.

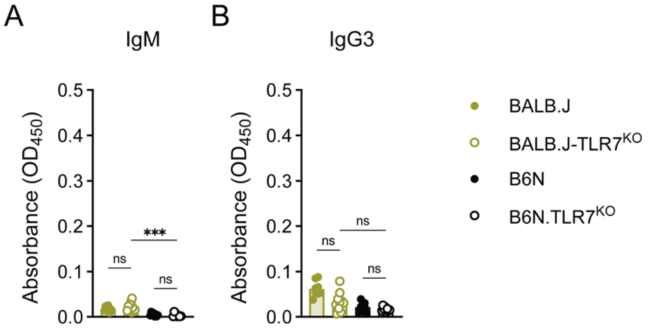


**Supplemental Figure 2. RL-MLV infection does not elicit an anti-IgM or -IgG3 antibody response.** (A) Sera from infected mice was collected 8 weeks post infection and monitored for IgM, (B) IgG3 against RL-MLV by enzyme-linked immunosorbent assay (ELISA). For each statistical comparison a nonparametric Kruskal-Wallis test was applied and corresponding significance values are indicated for each graph. ns, not significant; ***, $p<0.001$

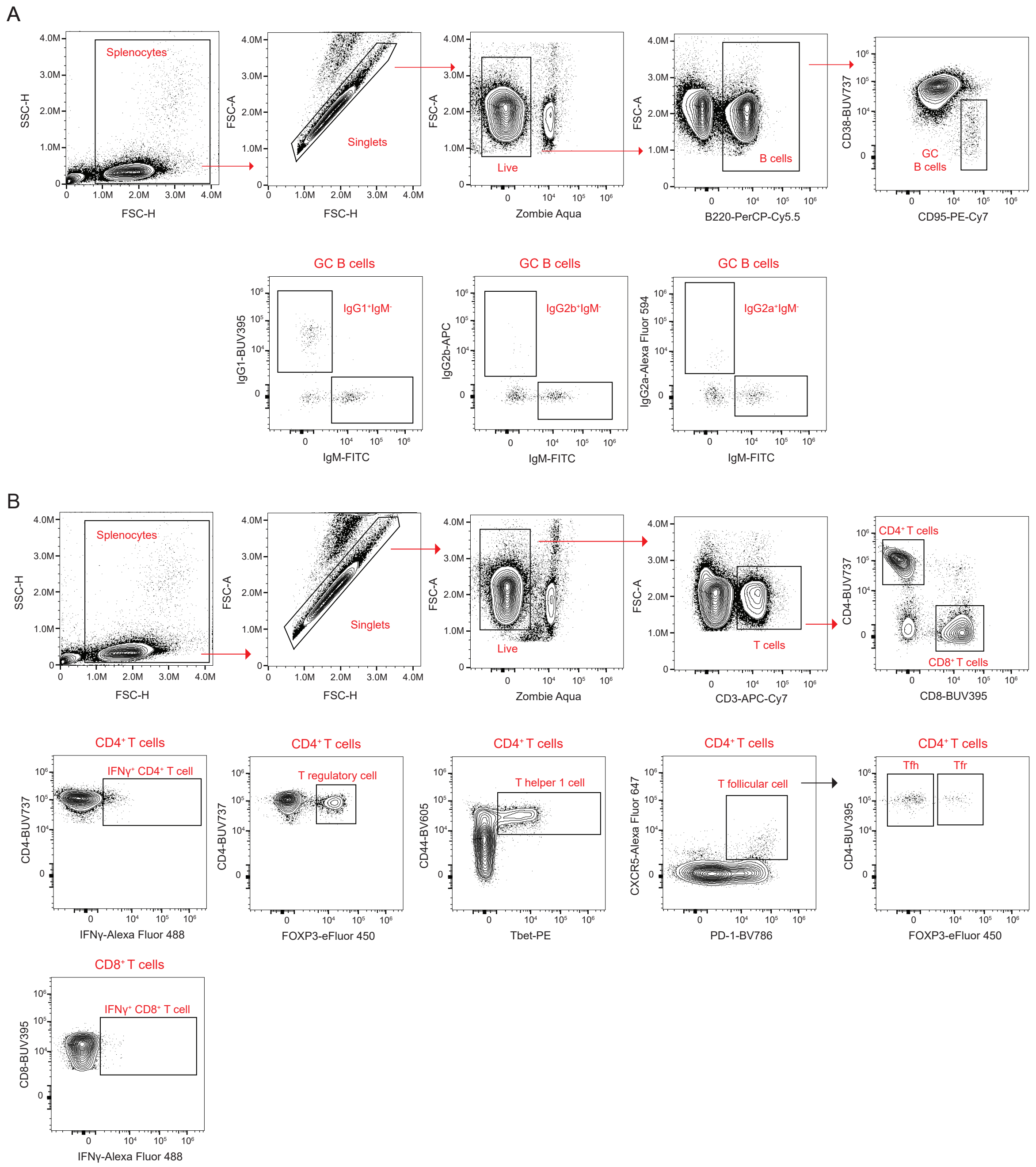


**Supplemental Figure 3. Gating Strategies** (A) Representative FACS plots of GC B cells in splenocytes. (B) Representative FACS plots of various CD4$^+$ T cell subsets in splenocytes.

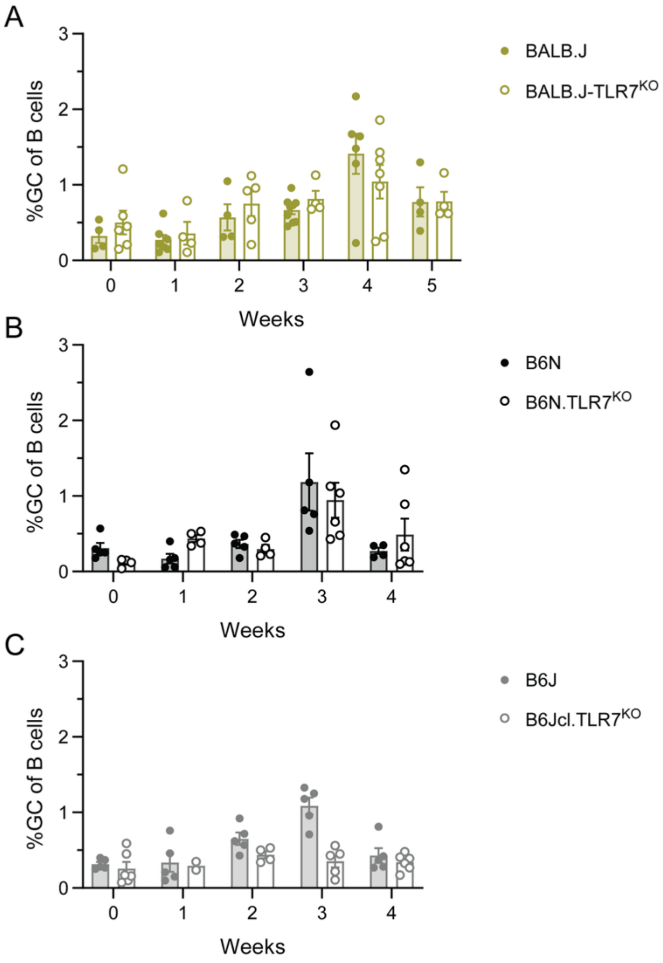


**Supplemental Figure 4. Upon infection with RL-MLV GC B cells peak 4 weeks post infection in BALB.J mice, and 3 weeks in B6J and B6N mice.** (A) BALB.J and TLR7-deficient BALB.J mice were infected with RL-MLV and monitored for frequency of GC B cells by FACS analysis weekly. (B) B6N and TLR7-deficient B6N mice were infected with RL-MLV and monitored for frequency of GC B cells by FACS analysis weekly. (C) B6J and TLR7-deficient B6Jcl mice were infected with RL-MLV and monitored for frequency of GC B cells by FACS analysis weekly.

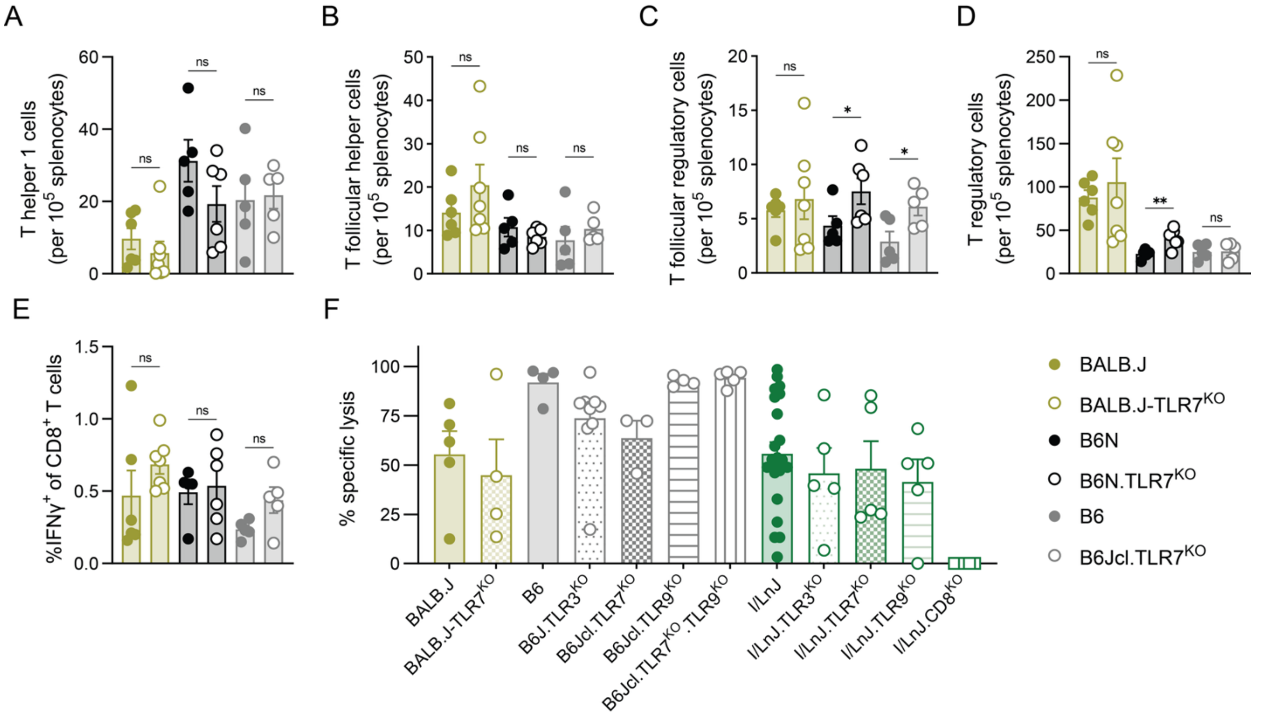


**Supplemental Figure 5. T cell responses in TLR7-deficient mice infected with MLV.** Mice of the indicated genotypes were infected with RL-MLV and splenocytes were isolated and analyzed by flow cytometry at peak GC response. (A) Number of T helper 1 cells (Live$^+$CD3$^+$CD4$^+$CD8$^-$CD44$^{hi}$T-bet$^+$). (B) Number of T follicular helper cells (Live$^+$CD3$^+$CD4$^+$CD8$^-$CXCR5$^+$PD-1$^+$FOXP3$^-$) (C) Number of T follicular regulatory cells (Live$^+$CD3$^+$CD4$^+$CD8$^-$CXCR5$^+$PD-1$^+$FOXP3$^+$) (D) Number of T regulatory cells (Live$^+$CD3$^+$CD4$^+$CD8$^-$FOXP3$^+$) (E) Frequency of IFNγ$^+$ CD8 T cells (Live$^+$CD3$^+$CD4$^-$CD8$^+$IFNγ$^+$) (F) Mice of the indicated genotypes were infected with RL-MLV. Donor splenocytes were incubated with either MLV GagL peptide, or DMSO. GagL loaded splenocytes were incubated with 5uM CFSE, while non-loaded splenocytes were incubated with 0.5uM CFSE. The CFSE-labeled populations are mixed at a 1:1 ratio and injected intravenously into infected mice. Splenocytes from infected mice were isolated 36 hours post injection and analyzed for CFSE expression by FACS and the frequency of specific lysis was calculated. *For each statistical comparison a nonparametric Kruskal-Wallis test was applied and corresponding significance values are indicated for each graph., not significant; *, p<0.05; **, p<0.01; ****, p<0.0001*

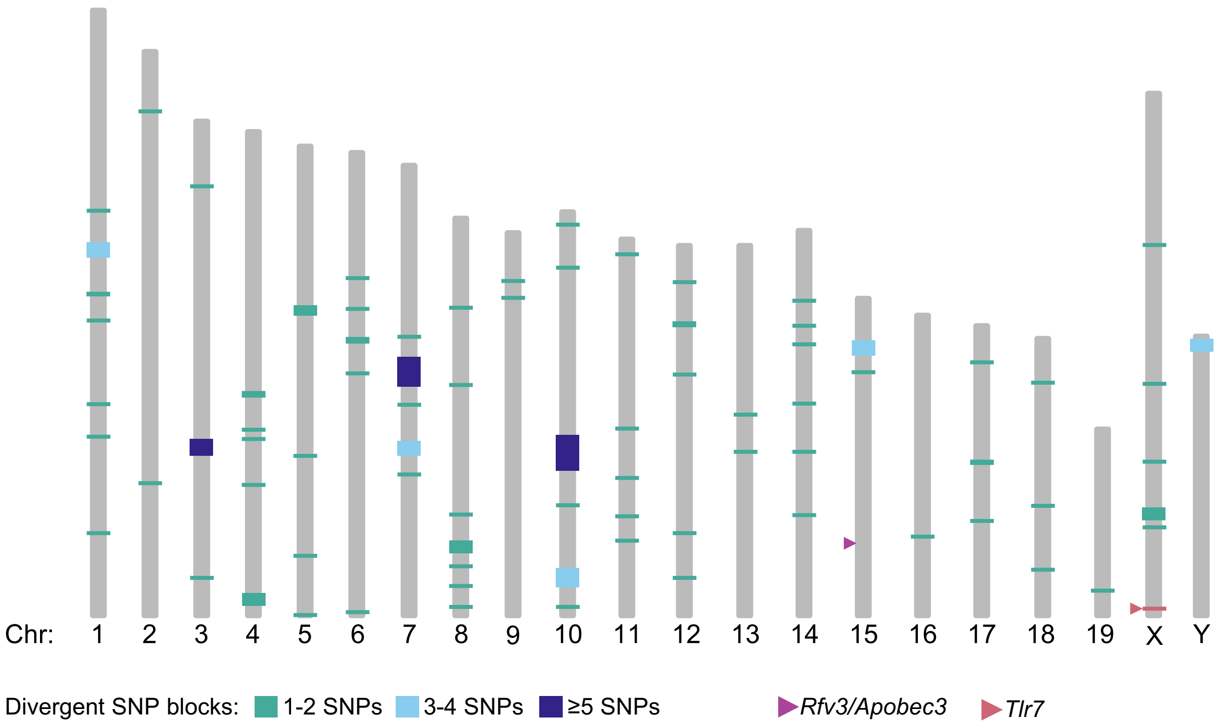


**Supplemental Figure 6. MiniMUGA comparison of B6N.TLR7$^{KO}$ and B6JcI.TLR7$^{KO}$ mice.** Diagram of chromosomes with locations of single nucleotide polymorphisms (SNPs) divergent between B6N.TLR7$^{KO}$ and B6JcI.TLR7$^{KO}$ mice as determined by MiniMUGA analysis of three mice from each line. SNPs less than 4Mb apart were grouped, and the number of SNPs in a block is color-coded as indicated. Locations of the MLV-resistance locus *Rfv3/Apobec3* and the *Tlr7* gene are indicated.

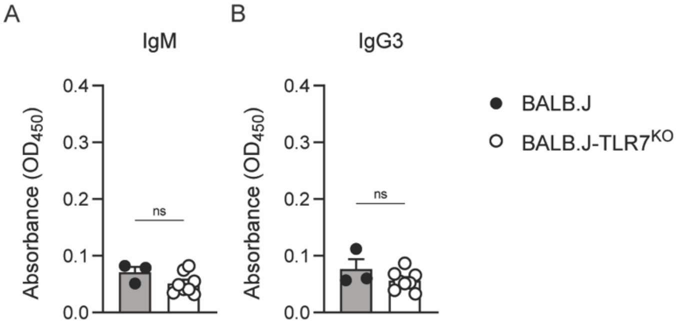


**Supplemental Figure 7. MMTV** BALB.J (closed circles) and TLR7-deficient BALB.J mice (open circles) were infected with MMTV. Sera from infected mice was collected 8 weeks post infection and monitored for (A) IgM and (B) IgG3 against MMTV virions by enzyme-linked immunosorbent assay (ELISA). *For each statistical comparison a nonparametric Kruskal-Wallis test was applied and corresponding significance values are indicated for each graph. ns, not significant.*

**Table S1. Antibodies and flow cytometry reagents utilized in this investigation. Antibodies are anti-mouse unless noted otherwise.**

| Antibody | Clone | Dilution | Manufacturer | Catalog number |
|---|---|---|---|---|
| Zombie Aqua™ Fixable Viability Kit | n/a | 1:1000 | BioLegend | 423101 |
| Zombie NIR™ Fixable Viability Kit | n/a | 1:1000 | BioLegend | 423106 |
| CD4-BUV737 | RM4-5 | 1:300 | BD Biosciences | 612844 |
| CD44-BV605 | IM7 | 1:300 | BD Biosciences | 563058 |
| CD279-BV786 | 29F.1A12 | 1:200 | BD Biosciences | 568566 |
| CD8a-BUV395 | 53-6.7 | 1:100 | BD Biosciences | 553027 |
| B220-PerCPCy5.5 | RA3-6B2 | 1:100 | BioLegend | 103236 |
| CD3-APCCy7 | 17A2 | 1:150 | BioLegend | 100222 |
| CXCR5-AF647 | L138D7 | 1:100 | BioLegend | 145532 |
| FOXP3-eFluor450 | FJK-16s | 1:100 | ThermoFisher | 48-5773-80 |
| IFNg-AF488 | XMG1.2 | 1:100 | BioLegend | 505815 |
| Tbet-PE | 4B10 | 1:100 | BD Biosciences | 561265 |
| CD38-BUV737 | 90/CD38 | 1:100 | BD Biosciences | 741748 |
| CD86-PE | PO3 | 1:100 | BioLegend | 105106 |
| CD95-PECy7 | JO2 | 1:100 | BD Biosciences | 557653 |
| CD138-BV786 | 281-2 | 1:100 | BD Biosciences | 569692 |
| IgG1-BUV395 | A85-1 | 1:300 | BD Biosciences | 740234 |
| IgG2b-APC | RMG2b-1 | 1:300 | BioLegend | 406711 |
| IgM-FITC | II/41 | 1:300 | BD Biosciences | 553437 |
| IgG2a-AF594 | RMG2a-62 | 1:300 | BioLegend | 407119 |
| CXCR4-BV421 | L276F12 | 1:200 | BioLegend | 146511 |
| CD4-PECy7 | GK1.5 | 1:500 | ThermoFisher | 25-0041-82 |
| VB6-PE | RR4-7 | 1:500 | BD Biosciences | 553194 |
| anti-STING | D2P2F | 1:1000 | CellSignaling | 13647 |
| anti-TUBA4A | B-5-1-2 | 1:10000 | Sigma-Aldrich | T6074 |
| goat anti-mouse IgG (H+L) HRP | n/a | 1:5000 (ELISA) 1:10000 (Immunoblot) | Jackson ImmunoResearch | 115-035-146 |
| goat anti-rabbit IgG (H+L) HRP | n/a | 1:5000 (ELISA) 1:10000 (Immunoblot) | Jackson ImmunoResearch | 111-035-144 |
| goat anti-mouse IgG2a HRP | n/a | 1:5000 | Jackson ImmunoResearch | 115-035-206 |
| goat anti-mouse IgG2c HRP | n/a | 1:2500 | Jackson ImmunoResearch | 115-035-208 |
| goat anti-mouse IgG1 HRP | n/a | 1:5000 | Jackson ImmunoResearch | 115-035-205 |
| goat anti-mouse IgG2b HRP | n/a | 1:5000 | Jackson ImmunoResearch | 115-035-207 |
| goat anti-mouse IgG3 HRP | n/a | 1:5000 | Jackson ImmunoResearch | 115-035-209 |
| goat anti-mouse IgM HRP | n/a | 1:5000 | Jackson ImmunoResearch | 115-035-075 |

**Supplementary Information References**
